# Cluster expansion of *apolipoprotein D (ApoD)* genes in teleost fishes

**DOI:** 10.1101/265538

**Authors:** Langyu Gu, Canwei Xia

## Abstract

**Background:** Gene and genome duplication play important roles in the evolution of gene function. Compared to individual duplicated genes, gene clusters attract particular attentions considering their frequent associations with innovation and adaptation. Here, we report for the first time the expansion of the ligand (e.g., pheromone and hormone)-transporter genes, *apolipoprotein D* (*ApoD*) genes in a cluster, specific to teleost fishes.

**Results:** The single *ApoD* gene in the ancestor expands in two clusters with a dynamic evolutionary pattern in teleost fishes. Based on comparative genomic and transcriptomic analyses, protein 3D structure comparison, evolutionary rate detection and breakpoint detection, orthologous genes show conserved expression patterns. Lineage-specific duplicated genes that are under positive selection evolved specific and even new expression profiles. Different duplicates show high tissue-specific expression patterns (e.g., skin, eye, anal fin pigmentation patterns, gonads, gills, spleen and lower pharyngeal jaw). Cluster analyses based on protein 3D structure comparisons, especially the four loops at the opening side, show segregation patterns with different duplicates. Duplicated *ApoD* genes are predicted to be associated with forkhead transcription factors and MAPK genes, and they are located next to the breakpoints of genome rearrangements.

**Conclusions:** Here, we report the expansion of *ApoD* genes specific to teleost fishes in a cluster manner for the first time. Neofunctionalization and subfunctionalization were observed at both protein and expression levels after duplication. Evidence from different aspects, i.e. abnormal expression induced disease in human, fish-specific expansion, predicted associations with forkhead transcription factors and MAPK genes, highly specific expression patterns in tissues related to sexual selection and adaptation, duplicated genes that are under positive selection, and their locations next to breakpoints of genome rearrangement, suggests the potential advantageous roles of *ApoD* genes in teleost fishes. Cluster expansion of *ApoD* genes specific to teleost fishes thus provides an ideal evo-devo model for studying gene duplication, cluster maintenance and new gene function emergence.

## Background

Gene and genome duplication play important roles in evolution by providing new genetic materials [1]. The gene copies emerging from duplication events (including whole genome duplications (WGD)) can undergo different evolutionary fates, and a number of models have been proposed as to what can happen after duplication [2]. In many instances, one of the duplicates becomes silenced via the accumulation of deleterious mutations (i.e. pseudogenization or nonfunctionalization [1]). Alternatively, the original pre-duplication function can be subdivided between duplicates (i.e. subfunctionalization) [3], or one of the duplicates can gain a new function (i.e. neofunctionalization) [4]. Since the probability of accumulating beneficial substitutions is relatively low, examples for neofunctionalization are sparse. There are, nevertheless, examples for neofunctionalization. For example, the duplication of *dachshund* in spiders and allies has been associated with the evolution of a novel leg segment [5]; the expansion of repetitive regions in a duplicated trypsinogen-like gene led to functional antifreeze glycoproteins in Antarctic notothenioid fish [6]; and the duplication of *opsin* genes is implicated in trichromatic vision in primates [7]. Another selective advantage of gene duplication can be attributed to the increased numbers of gene copies, e.g. in the form of gene dosage effects [8,9]. Multiple mechanisms can act together to shape different phases of gene evolution after duplication [10].

Gene functional changes after duplication can occur at the protein level [6,11,12]. For example, the physiological division of labour between the oxygen-carrier function of haemoglobin and the oxygen-storage function of myoglobin in vertebrates [13] (subfunctionalization); and the acquired enhanced digestive efficiencies of duplicated gene encoding of pancreatic ribonuclease in leaf monkeys [14] (neofunctionalization). However, the chance to accumulate beneficial alleles is low, and thus, functional changes after duplication at the protein level are sparse. Instead, changes in the expression level are more tolerable and efficient, since they do not require modification of coding sequences, and they can immediately offer phenotypic consequences. For example, complementary degenerative mutation in the regulatory regions of duplicated genes is a common mechanism for subfunctionalization [2,15]. Many examples have provided evidence that duplicated genes acquiring new expression domains are linked to neofunctionalization (e.g., *dac2*, a novel leg segment in Arachnid [5]; *elnb*, bulbus arteriosus in teleost fishes [16]; and *fhl2b*, egg-spots in cichlid fishes [17]).

In some cases, gene clusters resulting from gene duplication have attracted considerable attention, such as Hox gene clusters [18], globin gene clusters [19], paraHox gene clusters [20], MHC gene clusters [21] and opsin gene clusters [22]. Duplicated genes in clusters are usually related to innovation and adaptation [11,22,23], suggesting advantageous roles during evolution. The expansion of gene clusters can be traced back to WGD and tandem duplication [23,24]. Compared to two rounds of WGD occurring before the split between cartilaginous and bony fish [25], the ancestor of teleost fishes experienced another round of WGD (teleost genome duplication, TGD) after its divergence from non-teleost actinopterygians, including Bichir, Sturgeon, bowfin and spotted gar [26,27]. This extra TGD provides an additional opportunity for gene family expansion in fishes [22,28-30].

Genome rearrangements have been suggested to occur frequently in teleost fishes [31]. If genome rearrangements can capture locally adapted genes or antagonist sex determining genes by reducing recombination, the rearranged genome can promote divergence and reproductive isolation [32,33], and thus contribute to speciation and adaptation. Examples can be found in butterflies [34], mosquitoes [35] and fish [36]. This occurs especially when advantageous genes are located next to the breakpoints of a genome rearrangement due to its low recombination rates [37,38]. Considering that the expansion of gene clusters is usually adaptive (as mentioned above) and is linked to genome instability [33,39,40], it will be interesting to investigate the roles of gene clusters located next to the breakpoints of genome rearrangements. However, related studies are sparse.

Here, we report, for the first time, the expansion of the gene *apolipoprotein D* (*ApoD*) in teleost fishes. The *ApoD* gene belongs to the lipocalin superfamily of lipid transport proteins [41,42]. In humans, ApoD is suggested to function as a multi-ligand, multifunctional transporter (e.g., hormone and pheromone) [42,43], which is important in homeostasis and in the housekeeping of many organs [43]. It is expressed in multiple tissues, most notably in the brain and testis (see e.g. [42,44,45]), and it has been suggested to be involved in the central and peripheral nervous systems [42]. Interestingly, the *ApoD* gene is duplicated in a cluster manner in two chromosomes in different teleost fishes (http://www.genomicus.biologie.ens.fr/genomicus-91.01/cgi-bin/search.pl). However, no detailed analyses have been reported. Here, we investigated the evolutionary history of *ApoD* genes in fishes for the first time.

## Results

### 1. *In silico* screen and phylogenetic reconstruction of *ApoD* genes

To investigate the expansion of *ApoD* genes, we performed phylogenetic reconstruction with high quality assembly genome data (Figures 1A and 1B). There is one *ApoD* gene in coelacanth (*Latimeria chalumnae*), and two copies (*A* and *B*) in spotted gar (*Lepisosteus oculatus*), which are located in one cluster. Different numbers of *ApoD* genes are located in two clusters in different teleost fishes, i.e., with two copies in cavefish (*Astyanax mexicanus*) (*A1* and *A2*) and in pufferfish (*Tetraodon nigroviridis*) (*B2a* and *A2*); three copies (*A1, A2, B2*) in zebrafish (*Danio rerio*) and in cod (*Gadus morhua*) (*A1, A2* and *B2b*); four copies in platyfish (*Xiphophorus maculatus*) (*A1, A2, B2a* and *B2b*); five copies in Amazon molly (*Poecilia formosa*) (*A1, A2, B2a, B2ba1, B2ba2*) and in fugu (*Takifugu rubripes*) (*A1, A2, B1, B2a, B2b*); six copies in medaka (*Oryzias latipes*) (*A1, A2m1, A2m2, A2m3, B2a, B2b*); and eight copies in stickleback (*Gasterosteus aculeatus*) (*A1, A2s1, A2s2, B1, B2a, B2bs1, B2bs2, B2bs3*) and in tilapia (*Oreochromis niloticus*) (*A1, A2t1, A2t2, A2t3, A2t4, B1, B2a, B2b*). Noticeably, although the copy *B2b* of cod is clustered within the *B2a* clade based on a maximum likelihood (ML) tree, the bootstrap value is very low which could be due to recent duplication (Figure 1B). Considering its gene direction and syntenic with *B2b* genes in other species (Figure 1A), we named it copy *B2b*. Sequence alignment for tree construction can be found in Additional file 1.

**Figure 1.**
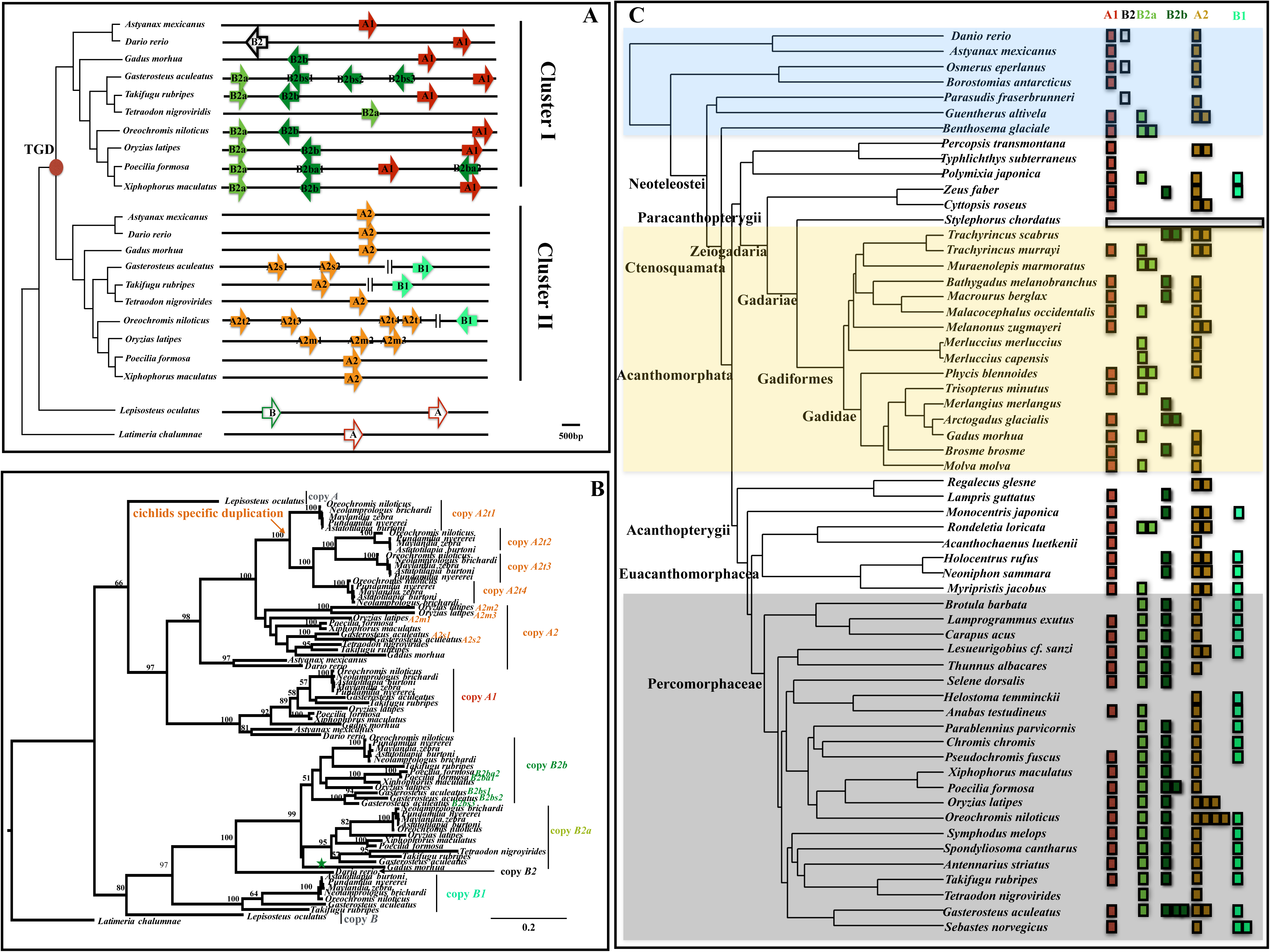
(A) Cluster expansion of *ApoD* genes specific to teleost fishes after teleost-specific duplication (TGD). Each arrow represents a single gene copy. Genes with red and orange colours represent paralogs derived from one common ancestor. Genes with dark and light green colours represent paralogs derived from one common ancestor. Phylogeny reconstruction is based on a consensus fish phylogeny [22]. (B) Maximum likelihood phylogenetic tree reconstruction to infer gene duplication. Bootstrap values > 50% are marked on the branch. Lineage-specific duplication events were labelled. Note, although copy *B2b* of *Gadus morhua* is clustered within the *B2a* clade (labelled with a green star), its bootstrap value is low, which could be due to recent duplication. (C) A dynamic evolutionary pattern of *ApoD* genes across the entire phylogeny of teleost fishes. Highly variable copy numbers are detected in different lineages, especially in the Paracanthopterygii lineage. *Stylephorus chordatus* lost all *ApoD* genes. Compared to copy *A1*, copy *A2* exhibits more variable lineage-specific duplicates in different fishes, with the highest numbers appearing in tilapia (four copies). Copy *B1* is absent in the whole clade of Gadiformes. Copy *B2* only shows up in species *Danio rerio, Osmerus eperlanus* and *Parasudis fraserbrunneri*. The co-existence of *B2a* and *B2b* is common in Percomorphaceae. The largest numbers of lineage-specific duplicated genes are found in tilapia (copy *A2*), medaka (copy *A2*) and stickleback (copy *B2b*) in Percomorphaceae.

To further retrieve the evolutionary history of *ApoD* genes in fishes, we performed an *in silico* screen across the whole phylogeny of teleost fishes using draft genomes (Figure 1C). No *ApoD* gene was detected in *Stylephorus chordatus*. Unlike copy *A1* which shows no lineage-specific duplication, copy *A2* exhibits variable lineage-specific copies in different fishes, with the highest lineage-specific duplication present in the cichlid fish tilapia (four copies). Copy *B1* is absent in the clade of Gadiformes but is kept in Acanthopterygii. Species in *Danio rerio, Osmerus eperlanus* and *Parasudis fraserbrunneri* possess copy *B2*. The co-existence of *B2a* and *B2b* is common in Percomorphaceae. The largest numbers of lineage-specific duplicated *ApoD* genes were found in tilapia (copy *A2*), medaka (copy *A2*) and stickleback (copy *B2b*) in Percomorphaceae. The predicted *ApoD* gene sequences can be found in Additional file 2.

To infer the relationship between *ApoD* gene duplication and TGD, syntenic analyses were conducted, revealing that two paralogous regions in teleost fishes correspond to one ohnolog region in spotted gar and in chicken (connected by the same coloured lines in Figure 2A). For example, the region with yellow colour in linkage group (LG) 9 in spotted gar (left above), or in chromosome (chr) 2 in chicken (left down) have paired paralogous regions (connected by yellow lines) in chr17 (*ApoD* cluster I located on) and in chr20 (*ApoD* cluster II located on) in medaka. Similarly, the region with purple colour in LG3 in spotted gar and in chr1 in chicken has paired paralogous regions (connected by purple lines) in chr17 and in chr20 in medaka.

**Figure 2.**
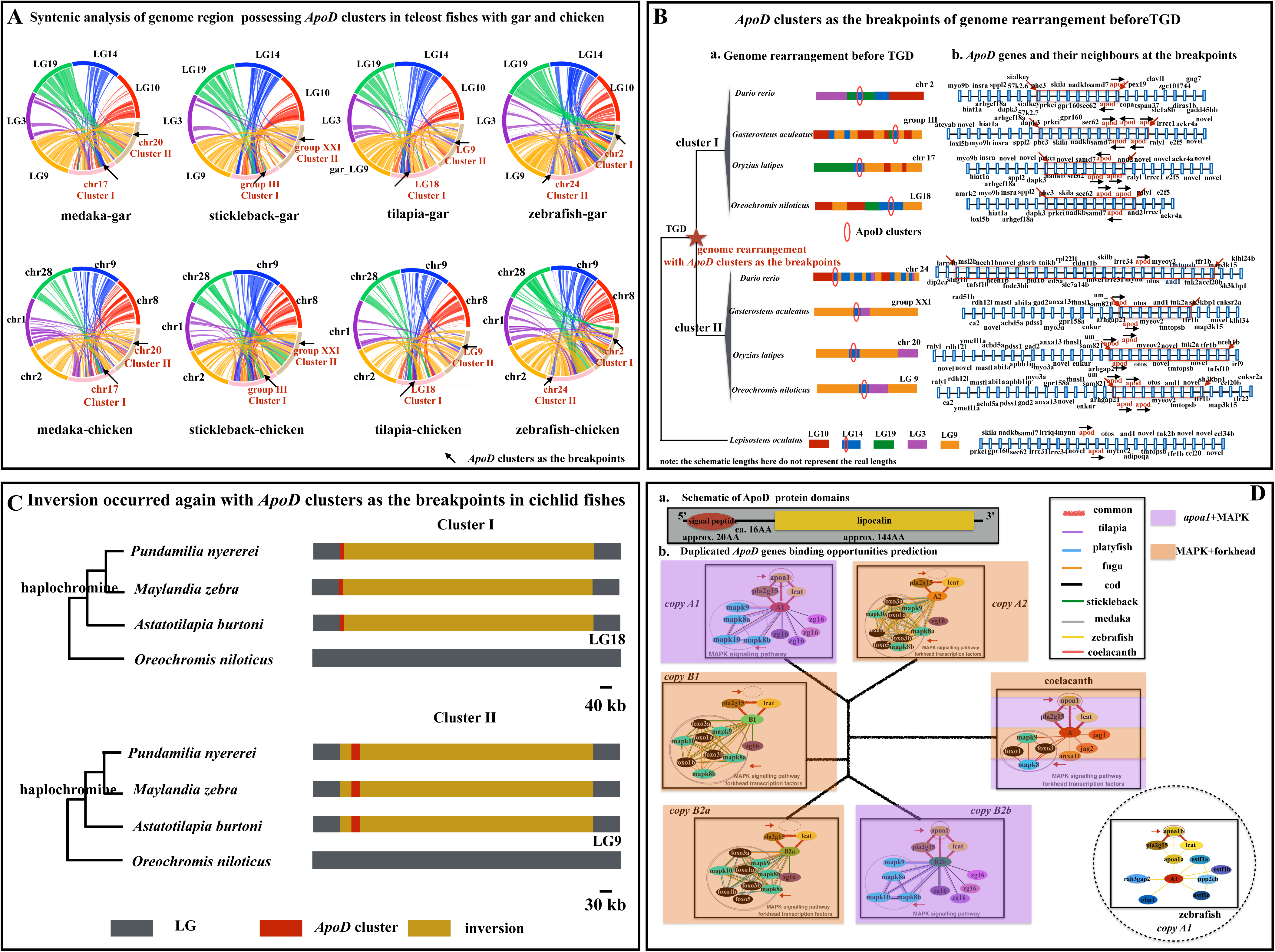
(A) Syntenic analyses between teleost fishes and spotted gar (top) and chicken (bottom), respectively. The same colour between spotted gar and chicken represents orthologous chromosomes. Two paralogous duplicated segments in teleost fishes (chromosomes (chrs)/linkage groups (LG) indicated in red colour) can be traced back to one corresponding orthologous region in spotted gar and chicken (chrs/LGs named with black colour), linked by colourful lines. Arrows show the regions where *apolipoprotein D* (*ApoD*) clusters are located. (B) *ApoD* clusters as the breakpoints of genome rearrangements before teleost-specific genome duplication (TGD). The same colour between gar and chicken represents orthologous chromosomes. Neighbouring genes are labelled. The red arrows show breakpoints, and the black arrows show gene direction. Neighbouring genes are named according to Ensembl database. (C) Inversion with *ApoD* clusters as the breakpoints occurred again in cichlid fishes. The haplochromine lineage, the most species-rich lineage of cichlid fishes, is labelled. (D) ApoD domains and protein-protein association predictions. a. Conserved domains include a single peptide of approx. 20 amino acids (AA) and a lipocalin domain of approx. 144 AA. b. Different paralogs exhibited differential associations. The common association is with MAPK genes. One class (copies *A2, B2a* and *B1*) is associated with multiple forkhead transcription factors. The other class (copies *A1* and *B2b*) lost this association; instead, it is associated with the lipoprotein-related gene *apoa1*. The single *ApoD* gene in coelacanth possesses both associations.

### 2. *ApoD* clusters next to the breakpoints of genome rearrangements

Syntenic analyses clearly show that more than one chromosome in spotted gar and in chicken are syntenic to the corresponding paralogous regions where *ApoD* clusters are located in fishes. For example, chromosomes (or LGs) containing *ApoD* clusters (chr17 and chr 20 in medaka, LG III and LG XXI in stickleback, LG9 and LG18 in tilapia, chr2 and chr24 in zebrafish, Figures 2A and 2B) are syntenic to chromosomes LG10, LG9, LG19, LG3 and LG14 in spotted gar and chr8, chr2, chr28, chr1 and chr9 in chicken (Figures 2A and 2B). Noticeably, *ApoD* clusters are located next to the breakpoints of genome rearrangements (Figures 2A and 2B). The rearranged segments (approx. 80 kb to 200 kb syntenic with LG14 in spotted gar, Figure 2B) include *ApoD* clusters and their neighbour genes (e.g., *and2, samd7, sec62, nadkb, gpr160, skila, prkci* and *phc3* for Cluster I; *otos, myeov2, and1, tmtopsb, tnk2a* and *tfr1b* for Cluster II) (Figure 2B). Analyses of available cichlid genomes further show that inversion of the segments (approx. 600 kb to 800kb) with *ApoD* clusters as the breakpoints also occurred in the species belonging to the most species-rich lineage, the haplochromine lineage in cichlid fishes, supported by split-read analysis using Delly (Figure 2C).

### 3. Protein-protein association predictions and protein 3D structure comparisons

To assess the biological functions of different *ApoD* genes, protein-protein associations were predicted. Protein domain architecture analyses revealed that ApoD proteins in different fishes are composed of approx. 20 amino acids as the signal peptide and approx. 144 amino acids as the lipocalin domain (Figure 2D). The common associations of *ApoD* genes are with the immunity-related gene *pla2g15* [47,48], the high-density lipoprotein biogenesis-related gene *lcat* [49], different copies of *zg16* related to pathogenic fungi recognition [50] and *MAPK* genes [51,52]. Different *ApoD* genes in teleost fishes can be subdivided into two classes based on the association predictions. *ApoD* genes from one class (copy *A2*, copy *B2a* and copy *B1*) are associated with forkhead transcription factors. *ApoD* genes from the other class (copy *A1* and copy *B2b*) lost their associations with forkhead transcription factors but are associated with the lipoprotein-related gene *apoa1* [53]. Noticeably, the only *ApoD* gene in coelacanth possesses both associations with forkhead transcription factors and with *apoa1*. *ApoD* in coelacanth is also associated with genes encoding ligands that can activate the Notch signalling pathway (*jag1, jag2*) [54], and the gene belongs to the annexin family (*anxaII*) (Figure 2D). Noticeably, more members of forkhead transcription factors and MAPK genes are associated with *ApoD* genes after duplication. Unlike in other fishes, copy *A1* in zebrafish is associated with bone resorption-related duplicated genes (*ostf1a, ostf1b, ostf1c*) [55], the cell growth and division-related gene *ppp2cb* [56], the neurodevelopment-related gene *rab3gap2* [57] and the guanylate-binding gene *gbp1* [58] (Figure 2D).

Homology protein structure modelling of different *ApoD* genes shows a conserved structure, including a cup-like central part made by eight antiparallel *β*-sheet strands and two ends connected by loops (a wide opening side formed by loops 1, 3, 5 and 7; and a narrow, closed bottom formed by loops 2, 4 and 6) (Figure 3A). Interestingly, unlike the very conserved cup-like central part, the loops are highly variable (Figure 3A). Cluster analyses based on the whole protein 3D structure comparison can clearly segregate different duplicates, e.g., copy *A1*, copy *A2*, copy *B2a* and copy *B2b*. Especially for lineage-specific duplicated genes which are clustered together. Copy *B1s* are clustered together, nesting within the copy *A2* clade (Figure 3B). The same analyses focused on only the four loops (loops 1, 3, 5 and 7) at the opening side show a similar segregation pattern (Figure 3C). However, cluster analyses focused on only the three loops (loops 2, 4 and 6) at the bottom side did not show a segregation pattern (Figure 3D). Details about the PDB files and the cluster results can be found in Additional file 3.

**Figure 3.**
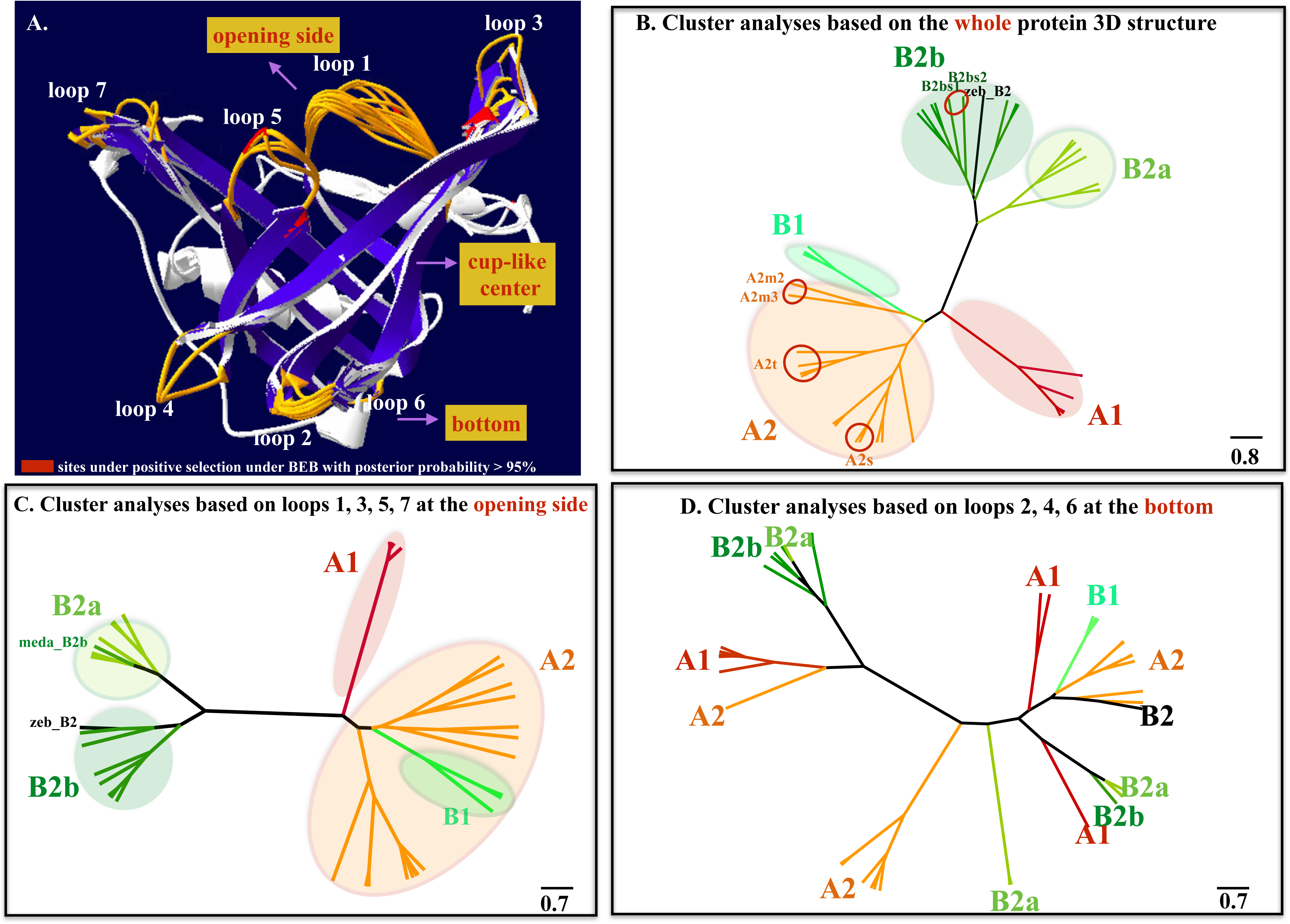
(A) Protein 3D structure modelling. Different ApoDs among species exhibited a conserved 3D structure, including a cup-like central part made by eight antiparallel *β*-sheet strands and two ends connected by loops (a wide opening part formed by loops 1, 3, 5 and 7; and a narrow, closed bottom formed by loops 2, 4 and 6). Unlike the very conserved cup-like central part, the loops at the two ends are highly variable. Note that most sites under positive selection are located on the loops or on the connections between the loops and the cup-like central part. (B) Cluster analyses based on the whole protein 3D structure. Cluster analyses can clearly segregate different *ApoD* duplicates, including lineage-specific duplicated genes. (C) Cluster analyses based on only four loops (loops 1, 3, 5 and 7) at the opening side. Cluster analyses can segregate different *ApoD* duplicates. (D) Cluster analyses based on only three loops (loops 2, 4 and 6) at the bottom side. Cluster analyses cannot segregate with different duplicates.

### 4. Positive selection detection on *ApoD* genes

Positive selection was detected in many *ApoD* genes before and after duplication, for example, the branch of copy *A2* in zebrafish, the ancestral branch of copies *B2a* and *B2b* in fugu, medaka and platyfish; and, especially, in lineage-specific duplicated genes, such as in stickleback, cichlid fishes, and amazon molly (Figure 4 and Additional file 4). Positive selection sites under Bayes empirical Bayes (BEB) with posterior probability >95% were given in Figure 5 and Additional file 4. Noticeably, most sites under positive selection are located on the highly variable loops or on the connections between loops and the cup-like central part at the opening side (loops 3 and 5) (Figures 3A and 5). Our strategy here is relatively conservative. As we can see, if we only focus on one prior hypothesis on a particular branch without doing multiple test correction, we will find more branches with dN/dS that are significantly larger than 1 (*p*<0.05) (Additional file 4). However, we chose to use our strategy to make the analyses more rigorous. Even with this strict but comprehensive strategy, our results still show that positive selection occurred in branches before and after *ApoD* gene duplication, especially for lineage-specific duplicated genes. Details about the codon alignment for the PAML analyses can be found in Additional file 4.

**Figure 4.**
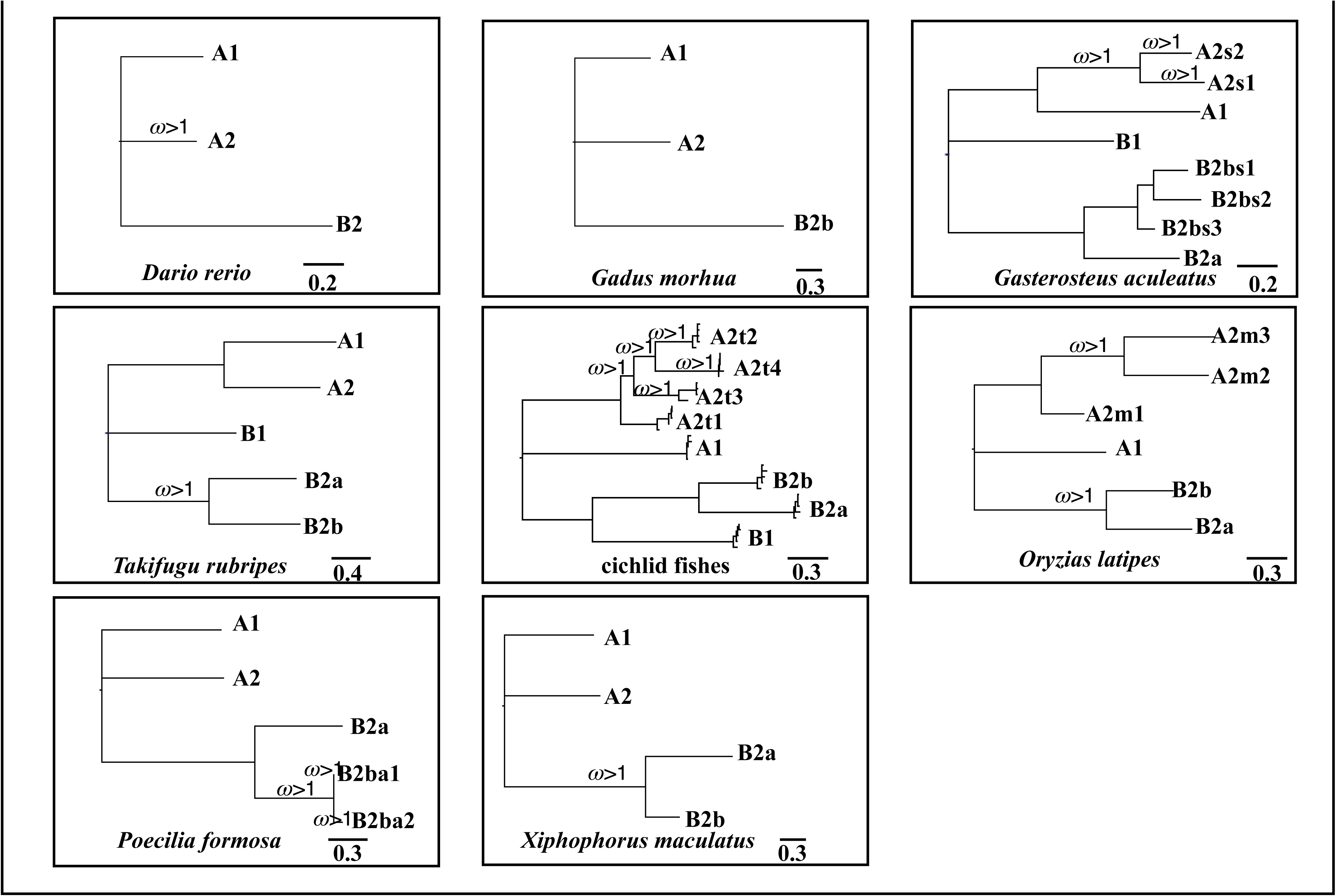
Positive selection detection of *ApoD* genes in different fishes under the branch-site model. Many duplicated *ApoD* genes are under positive selection with the value of ω significantly larger than 1 after Bonferroni correction, especially for lineage-specific duplicated genes. Note that if the whole clade is designated as the foreground branch when detecting positive selection, each branch of this clade will be labelled if its ω is significantly larger than 1. The unrooted tree was used for PAML analyses, and the rooted tree is only for presentation. More details can be found in Additional file 4.

**Figure 5.**
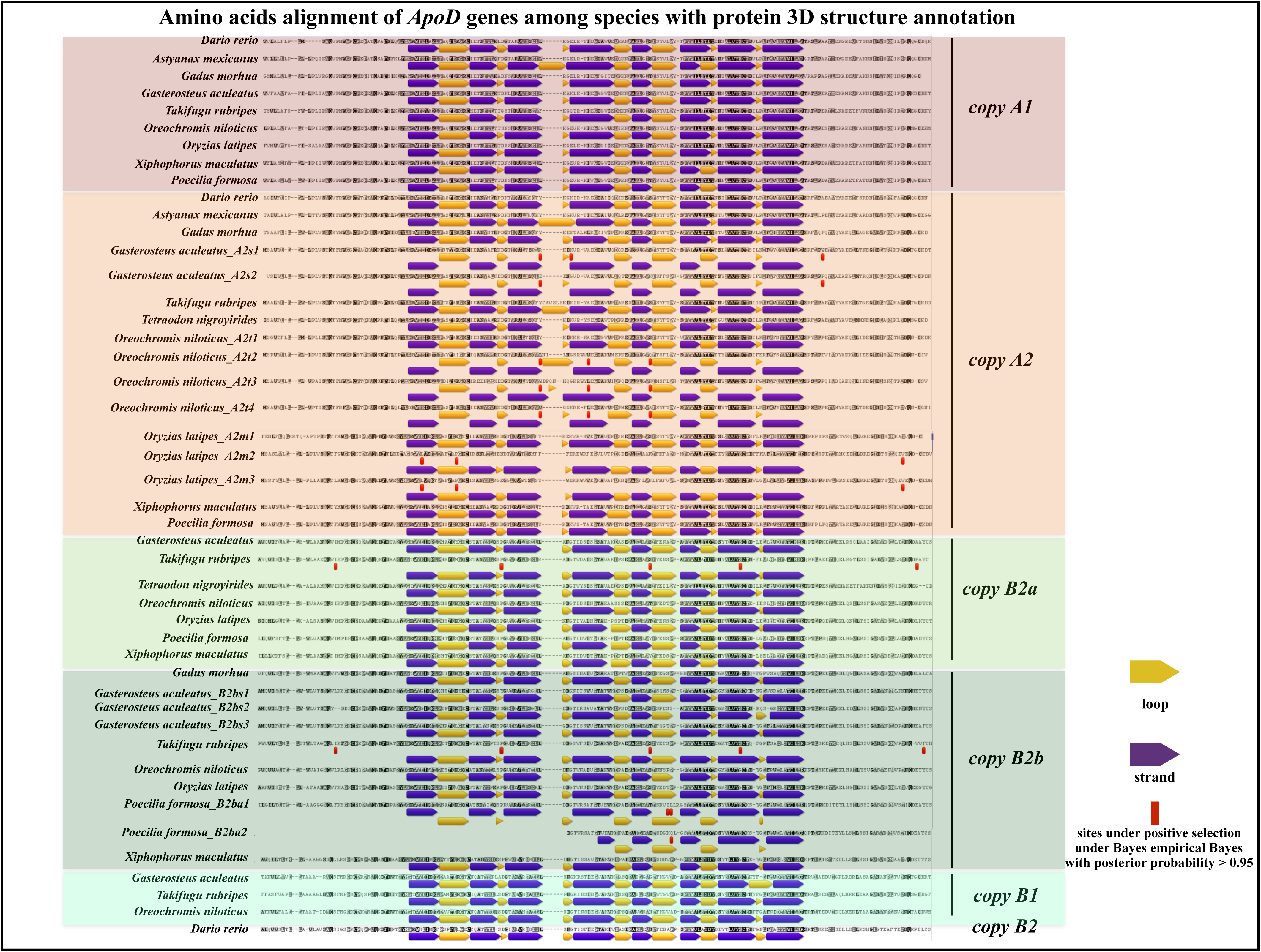
Amino acids alignment of *ApoD* duplicates among species with annotation. The purple arrow represents the *β*-strand, and the yellow arrow represents the loop. Sites under positive selection under Bayes empirical Bayes (BEB) with posterior probability > 95% are labelled with the red colour. Most sites under positive selection are distributed on the loops or on the connections between the loop and the *β*-strand. More details can be found in Additional file 4. Note, the positions of the sites that under positive selection in Additional file 4 are based on the codon alignment used for PAML analyses (Additional file 4) instead of the amino acids alignment here. But the amino acids that under positive selection are the same in Figure 5 and Additional file 4.

### 5. Gene expression profile detection of *ApoD* genes in different species

Copy *A1* is mainly expressed in the skin and the eye. Noticeably, copy *A1* is also highly expressed in the novel anal fin pigmentation patterns of a cichlid fish, *Astatotilapia burtoni*, which is in consistent with our previous study [59]. Copy *A2* shows redundant expression patterns in the gills, eye, skin and gonads in zebrafish and cavefish, but its orthologous *A2* genes including lineage-specific duplicated ones in the species of Acanthomorphata, which we test here (cod, stickleback, medaka, tilapia and *A. burtoni*), are expressed in the gills and in the related apparatus, the lower pharyngeal jaw. Copy *B2* in zebrafish shows redundant expression profiles (in the skin and gonads) overlapping with the profiles of copies *A1* and *A2*. Copies *B2a* and *B2b* show specific expression patterns in different fishes of Acanthomorphata, which we test here. For example, copy *B2b* is expressed in the ovary and in the liver in cod, while *B2a* and *B2bs* are mainly expressed in the spleen and in the liver of the stickleback. The expression of *B2a* and *B2b* was not detected in adult medaka, tilapia or *A. burtoni*. Interestingly, based on the available transcriptomic data for different developmental stages of gonads in tilapia, we found that *B2a* and *B2b* are highly specifically expressed at the early developmental stage of gonad tissues (five days after hatching, 5dah) (Figure 6). Copy *B1* is highly specifically expressed in the liver in tilapia and in *A. burtoni* but not in stickleback in which *B1* is inverted (Figure 1A). Noticeably, no expression profile was detected for copy *B* in gar, at least in the tissues we test here. Instead, expression profiles including the skin, eye, gill, liver, testis and brain were detected for copy *A* in gar. Details can be found in Figure 6 and Additional file 5.

**Figure 6.**
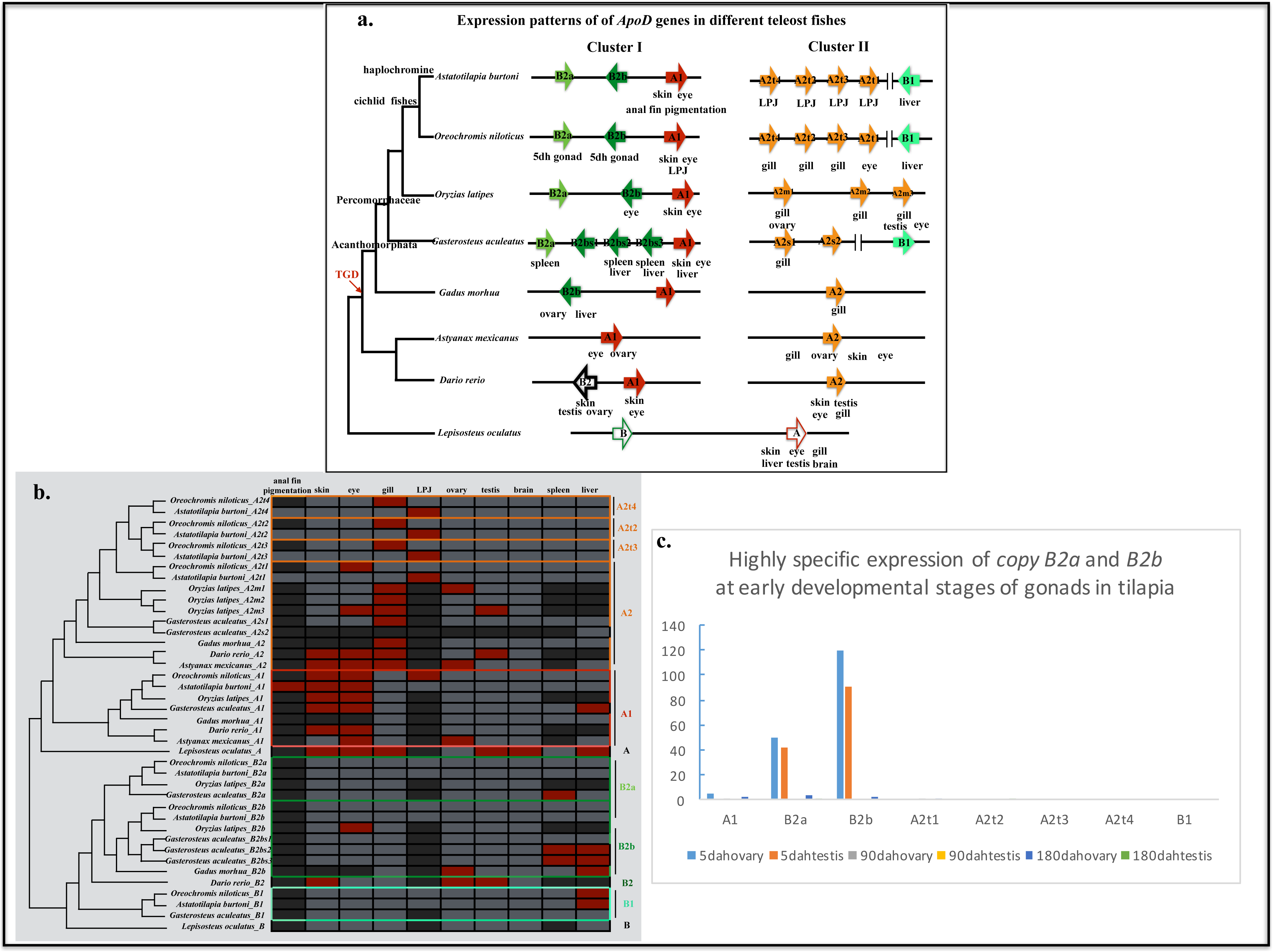
Gene expression profiles of different *ApoD* genes. (a) Schematic figure showing the gene expression profiles of different *ApoD* genes in two clusters after teleost-specific duplication (TGD). Each block represents a single gene copy. Phylogeny reconstruction was based on the consensus fish phylogeny [22]. (b) Detailed gene expression profiles of *ApoD* genes among fishes. The red colour represents tissues with a high expression level. Dark grey, data unavailable (either because the tissue does not exist in the species, or is undetected in this study). Light grey represents tissues with a low expression level. LPJ, the lower pharyngeal jaw in cichlid fishes. (c) Gene expression profiles of *ApoD* genes in different developmental stages of gonads in tilapia based on available transcriptomic data [115]. RPKM, reads per kilobase per million mapped reads; dah, days after hatching.

## Discussion

### Dynamic evolution of *ApoD* genes in teleost fishes

Here, we report the cluster expansion of *ApoD* genes specific to teleost fishes for the first time. Phylogenetic reconstruction and syntenic analyses clearly show the expansion of *ApoD* genes in two clusters with lineage-specific tandem duplications after TGD. The interplay between genome duplication and tandem duplication to prompt the expansion of the gene family has already been shown in a few studies [60-62]. It has been suggested that the fixation of duplications is much more common in genome regions where the rates of mutations are elevated due to the presence of already-fixed duplication, which is the so-called “snow-ball” effect [63]. Both genome duplication and tandem duplication can produce raw genetic materials to fuel the diversification of teleost fishes at a later stage, e.g., morphological complexity and ecological niche expansion, which is compatible with the “*time lag model*” [64]. Neofunctionalization and subfunctionalization after duplication are important steps to realize this diversification, and both can be detected in *ApoD* duplicates.

Functional divergence of *ApoD* genes occurred at the protein level. On the one hand, based on protein-protein association predictions, although common associations with MAPK genes were shared among *ApoD* genes, subdivided associations with forkhead transcription factors and *apoa1* were detected in different paralogous *ApoD* genes, indicating subfunctionalization. Noticeably, more members of forkhead transcription factors and MAPK genes are associated with *ApoD* genes after duplication. In this case, neofunctionalization could also occur but is not limited to the dosage effects. On the other hand, divergence was also detected at the protein structure level. Protein structures are often related to functional divergence [65-68]. New fold can even evolve novel function [69,70]. Even a few residues mutations can induce structural changes [71,72]. Therefore, structure-based inference is important for understanding molecular function. Our protein 3D structure modelling showed a conserved backbone conformation in spite of sequence divergence. Indeed, this is a feature of the lipocalin family [66], to which ApoDs belong [41,42,73]. The cup-like central part is used to transport large varieties of ligands, such as hormones and pheromones [73]. Interestingly, cluster analyses based on the whole protein structure or only on the four loops at the opening side can segregate different duplicates, although not for copy *B1.* Considering that copy *B1* has been lost multiple times, its function might be not necessary and could share functions with other copies; thus, it is not surprising if they are clustered together within the copy *A2* clade. The cluster results of two other individual genes (copy *B2* of zebrafish and *B2b* of medaka) do not affect the general segregation pattern. Noticeably, unlike the four loops at the opending side, cluster analyses based on only the three loops (loops 2, 4 and 6) at the bottom side did not show a clear segregation pattern (Figure 3D). It has been suggested that the loops of lipocalin proteins can affect ligand binding specificities, which are similar to the binding modes of antibodies [74]; and mediate protein-protein interactions [66]. Their segregation with different duplicates might indicate functional divergence occurred at the loops at the opening side. The evidence that most amino acids under positive selection are located on the loops or the connections between loops and strands (Figures 3A and 5, Additonal file 4) further indicates the potential important roles of these loops during evolution. Actually, reshaping different parts of the protein structure is an efficient way to produce functional divergence at the protein level in a short time [73], and this can be one of the ways in which divergence at the protein level occurred in *ApoD* genes.

Functional divergence of *ApoD* genes also occurred at the expression level. Different *ApoD* genes in zebrafish and cavefish exhibited redundant expression profiles but became more specific as the numbers of tandem duplicates increased, indicating subfunctionalization. Noticeably, new expression profiles were detected in duplicated paralogs, e.g., copy *A1* in novel anal fin pigmentation patterns and copy *A2s* in the lower pharyngeal jaw in *A. burtoni*, which is a representative cichlid fish of the most species-richness haplochromine lineage. These two traits are key innovations that are associated with adaptive radiation in cichlid fishes [75]. With expression changes, duplication can be an important source for the emergence of novelty [76], especially if they are adaptive. The relaxed selection pressure induced by the changing expression profiles can even further prompt the accumulation of mutations at the protein level.

### Potential advantageous roles of *ApoD* genes in teleost fishes

The *ApoD* gene has been thoroughly studied in humans and mice, and its abnormal expression was reported to be related to human diseases, such as Parkinson’s disease and Alzheimer’s disease [77-79]. Interestingly, as our study revealed here, *ApoD* dynamically expanded in fishes in two clusters in different chromosomes. Furthermore, these expanded duplicates are transcriptionally active. For example, they are highly expressed, and are restricted to different tissues related to sexual selection (skin [80], eye [81,82], gonads, anal fin pigmentation pattern [59]) and adaptation (gills [83-86], spleen [87,88], lower pharyngeal jaw [75,89]). These tissues are also derived from the neural crest, which is a key innovation in vertebrates [75,90-92]. Besides, many duplicated *ApoD* genes are under positive selection, especially for lineage-specific duplicated genes. Combined with their associations with MAPK genes and forkhead transcription factors, and their functions as pheromone and hormone transporters, we report evidences that suggests the potential importance of *ApoD* genes have in fishes.

Noticeably, two clusters exhibited distinct expression patterns. *ApoD* genes in cluster I in most fishes are expressed in tissues related to sexual selection (skin, eye, gonad and anal fin pigmentation), but in cluster II, *ApoD* genes are mainly expressed in tissues related to adaptation (gills and lower pharyngeal jaw). Interestingly, both clusters were maintained with their neighbouring genes and are located next to the breakpoints of genome rearangements during evolution. An inversion with *ApoD* clusters next to the breakpoints occurred again in cichlid fishes belonging to the haplochromine lineage, which is the most species-rich lineage [93]. If genome rearrangements can capture locally adaptive genes, or genes related to sexual antagonism, it would accelerate divergence by reducing recombination rates, thus prompting speciation and adaptation, even the emergence of neo-sex chromosome [94]. Given that many *ApoD* genes are under positive selection, their location at the breakpoints of genome rearrangements gives them more opportunities to be involved in speciation and adaptation, which deserves further investigation.

## Conclusions

Here, we report the cluster expansion of *ApoD* genes in a cluster manner specific to fishes for the first time. Different evidences based on computational evolutionary analyses strongly suggest the potentially advantageous roles of *ApoD* genes in fishes. An in-depth functional characterization of *ApoD* genes could help consolidate a model for the study of subfunctionalization and neofunctionalization. Moreover, finding the regulatory mechanisms behind the cluster expansion of *ApoD* genes and the reason why the *ApoD* gene expanded specific to fishes remain open questions. As more fish genomes with high assembly quality are available, especially closely related species such as cichlid fishes, the roles of *ApoD* genes in fish speciation and adaptation could be further investigated. Above all, the *ApoD* gene clusters reported here, provide an ideal evo-devo model for studying gene duplication, cluster maintenance, and the gene regulatory mechanism and their roles in speciation and adaptation.

## Methods

### *In silico* screen and phylogenetic reconstruction to infer gene duplication

To retrieve *ApoD* gene duplication in teleost fishes, we first extracted orthologs and paralogs in fishes with available genome data from Ensembl (Release 84) [95] and the NCBI database (https://www.ncbi.nlm.nih.gov/genome/). To confirm gene copy numbers, all orthologs and paralogs were used as queries in a tblastx search against the corresponding genomes. For all unannotated positive hits, a region spanning approx. 2 kb was extracted, and open reading frames (ORF) were predicted using Augustus (http://augustus.gobics.de/) [96]. The predicted coding sequences were then BLAST to the existing transcriptome database to retrieve the corresponding cDNA sequences. The cDNAs were then re-mapped to the corresponding predicted genes to recheck the predicted exon-intron boundary. Coding sequence of genes from human and spotted gar, as well as their neighbour genes were used to perform tblastx against the genomes of lamprey (*Lampetra fluviatilis*) and amphioxus (*Branchiostoma belcheri*). To infer gene duplication, a ML analysis was performed with *ApoD* genes retrieved from available assembled genomes in RAxML v8.2.10 [97] with the GTR+G model and ‘-f a’ option (which generates the optimal tree and conducts 10,000 rapid bootstrap searches).

To further retrieve *ApoD*s in other fishes across the whole phylogeny with draft genomes [21], all sequences retrieved above were used as queries in a tblastx search with the threshold *e* value 0.001. The hit scaffolds were retrieved, and genes within scaffolds were predicted using Augustus. The predicted *ApoD*s were then translated and re-aligned with known *ApoD* ORF to further confirm the exon-intron boundary. All predicted orthologs and paralogs were used as a query again in a tblastx search against the corresponding genome data until no more *ApoD* genes were predicted.

### Syntenic analyses and inversion detection

To further confirm the relationship between *ApoD* gene duplication and TGD, genes regions adjacent to the duplicates and to the outgroup species that did not experience TGD (including spotted gar and chicken) were retrieved. To this end, a window of 5 Mb around the *ApoD* clusters in teleost fishes as well as the corresponding chromosomes in spotted gar and chicken were retrieved from the Ensemble database and NCBI database. Syntenic analyses were performed with SyMap [98]. To further detect the structural variation around *ApoD* genes in cichlid fishes, we retrieved the available cichlid genomes raw data from [99]. The program Delly [100] based on paired-end split-reads analyses was used to detect the inversion and its corresponding breakpoints of the segments that contain the *ApoD* clusters in cichlid fishes, with tilapia as the reference.

### Protein-protein interaction prediction, protein 3D structure modelling and comparison

To predict the biological functions of *ApoD* genes, the protein domains of *ApoD* genes and protein-protein interactions were predicted with the Simple Modular Architecture Research Tool (SMART) [101,102] and the stringdb database http://string-db.org/ [103]. To see whether there are divergences at the protein level for different ApoDs, protein 3D structures were simulated with Swiss-model [104-106] using the human ApoD crystal protein structure (PDB ID 2hzr) as the template. The results were further visualized, evaluated and analysed with Swiss-PdbViewer [107]. To compare the protein structures, we first extracted and converted the corresponding information in the PDB file using Swiss-PdbViewer. Protein 3D structure comparisons were then conducted using Vorolign [108], which can compare closely related protein structures even when structurally flexible regions exist [108]. The protein 3D structure comparisons and cluster analyes were conducted with the ProCKSI-server (http://www.procksi.net/). Clustering results were further visualized using Figtree v1.4.2 (http://tree.bio.ed.ac.uk/software/figtree). Copy *B2ba2* of of amazon molly was not included in the cluster analyses due to its relatively short sequence. The PDB files were used as the input files. For the global protein structure comparisons, PDB files were extracted directly from the simulation results of Swiss-model [104-106]. To only get PDB files of the loop region, we revised the PDB files manually to get rid of the cup-like central part. To consider the highly variable connections between the loops and the cup-like central part (Figure 3A), we also included the structures of two more residues next to the loops. The resulting PDB files were further checked using Swiss-PdbViewer.

### Positive selection detection

To examine whether *ApoD* duplicates underwent adaptive sequence evolution, a branch-site model was used to test positive selection affecting a few sites along the target lineages (called foreground branches) in codeml within PAML [109,110]. The rates of non-synonymous to synonymous substitutions (ω or dN/dS) with a priori partitions for foreground branches (PAML manual) were calculated. Considering the very dynamic evolutionary pattern of *ApoD* genes, we tested positive selection in every branch in each species. In this case, we designated each branch as the foreground branch to run the branch-site model multiple times. Noticeably, there are two different ways to designate the foreground branch in the branch-site model. One way is to designate only the branch as the foreground branch. The other way is to designate the whole clade (all the branches within the clade including the ancestral branch) as the foreground branch. We included both perspectives in our data analyses (Additonal file 4). If the whole clade is under positive selection, we will label all the branches within this clade including the ancestral branch with ω>1 in the results (Figure 4).

All model comparisons in PAML were performed with fixed branch lengths (fix_blength = 2) derived under the M0 model in PAML. Alignment gaps and ambiguity characters were removed (Cleandata = 1). A likelihood ratio test was used to test for statistical significance. In addition, Bonferroni correction was conducted for multiple test correction [111]. The BEB was used to identify sites that are under positive selection. Sites under positive selection under BEB with posterior probability >0.95 are given in Figure 5 and Additional file 4.

### Gene expression profile analyses

To see the expression profiles of *ApoD* duplicates in different fishes, raw transcriptomic data from zebrafish, cavefish, cod, medaka, tilapia and *A. burtoni* were retrieved from NCBI https://www.ncbi.nlm.nih.gov/ (Additional file 5). Raw reads were mapped to the corresponding cDNAs (http://www.ensembl.org/) to calculate the RPKM (reads per kilobase per million mapped reads) value. Quantitative polymerase chain reaction (qPCR) was used to detect the expression profiles of different *ApoD* duplicates in the tissues that are not included in the existing transcriptomes (Additional file 5). Prior to tissue dissection, specimens were euthanized with MS 222 (Sigma-Aldrich, USA) following an approved procedure (permit nr. 2317 issued by the cantonal veterinary office, Switzerland; Guidelines for the Care and Use of Laboratory Animals prescribed by the Regulation of Animal Experimentation of Chongqing, China). RNA isolation was performed according to the TRIzol protocol (Invitrogen, USA). DNase treatment was performed with the DNA-free™ Kit (Ambion, Life Technologies, USA). The RNA quantity and quality were determined with a NanoDrop1000 spectrophotometer (Thermo Scientific, USA). cDNA was produced using the High Capacity RNA-to-cDNA Kit (Applied Biosystems, USA). Housekeeping gene elongation factor 1 alpha *(elfa*1) [112], ubiquitin (*ubc*) [113] and ribosomal protein L7 (*rpl7*) [17] [114] were used as endogenous control. qPCR was performed on a StepOnePlusTM Real-Time PCR system (Applied Biosystems, Life Technologies) using the SYBR Green Master Mix (Roche, Switzerland) with an annealing temperature of 58°C and following the manufacturer’s protocols. Primers are available in Additional file 6.

### List of abbreviations

ApoD: apolipoprotein D. WGD: whole genome duplication. TGD: teleost genome duplication. ML: maximum likelihood. LG: linkage group. chr: chromosome. BEB: Bayes empirical Bayes. ORF: open reading frame. SMART: Simple Modular Architecture Research Tool. RPKM: Reads Per Kilobase per Million mapped reads. qPCR: quantitative polymerase chain reaction. *elfa*1: elongation factor 1 alpha. *ubc:* ubiquitin. *rpl7:* ribosomal protein L7.

## Declarations

### Ethics approval and consent to participate

Animal experiments reported in this study have been approved by the cantonal veterinary office in Basel under permit number 2317, Switzerland; Guidelines for Care and Use of Laboratory Animals prescribed by the Regulation of Animal Experimentation of Chongqing, China.

### Consent for publication

Not applicable

### Availability of data and material

The datasets supporting the results of this article are available in the Dryad repository, doi:10.5061/dryad.39g63v2. https://datadryad.org/review?doi=doi:10.5061/dryad.39g63v2 [46].

## Competing interests

The authors declare that they have no competing interests.

### Funding

This work was supported by the PhD grant from University of Basel, Switzerland; postdoctoral funding from Southwest University, China; China Postdoctoral Science Foundation Grant (2018M633311); and National Natural Science Foundation of China (31802283) to LG.

### Author’s contributions

LG discovered the expansion of *ApoD* gene family, did data analysis and wrote the manuscript. LG did protein 3D structure comparison and cluster analyses with the help of CX.

## Acknowledgements

We thank Prof. Walter Salzburger for his suggestions. We also thank Prof. Deshou Wang for his help of sampling of tilapia and laboratory support. We particularly thank Prof. Ziheng Yang for his valuable suggestions about positive selection detection using PAML. Many thanks to the kind computing support from Prof. Zhengwang Zhang. We appreciate the valuable suggestions and comments from the associate editor and two anonymous reviewers very much. Many thanks to Dario Moser, Heinz-Georg Belting, Jing Wei and Hua Ruan for their help of fish sampling. Many thanks for the help of fish dissection from Yang Zhao, Fabrizia Ronco, Attila Rüegg, Adrian Indermaur, Xianbo Zhang and He Ma. Many thanks to Lukas Zimmerman, Peter Fields, Yuchen Sun, Yanyan Xu, Zihui Zhang and De Chen for their help and fruitful discussions. We also would like to express our thanks to Prof. Xiangjiang Zhan in the Institute of Zoology, Chinese Academy of Sciences for the help of revising the rebuttal letter. Many thanks to Prof. Anders Moller in CNRS, Université Paris-Sud in France, Dr. Juan Felipe Ortiz in Vanderbilt University in America for the correction of English language.

## Additional Files

**Additional file 1** Sequence alignment for ML tree reconstruction.

**Additional file 2** Predicted ApoD genes across the phylogeny.

**Additional file 3** Protein PDB files and cluster analyses results.

**Additional file 4** Sequence alignment for PAML test and statistics results.

**Additional file 5** Raw transcriptomic data information and gene expression profiles.

**Additional file 6** qPCR primer sequences.

